# Flavin-based extracellular electron transfer under oxic conditions by Gram-positive *Microbacterium deferre* sp. nov. A1-JK in redox-graded environments

**DOI:** 10.1101/2024.09.12.612730

**Authors:** Jamie J.M. Lustermans, Naja Basu, Leonid Digel, Kartik Aiyer

**Affiliations:** Microbial Systems Technology Excellence Centre, University of Antwerp, Belgium; Department of Biology, Research Group Geobiology, University of Antwerp, Wilrijk, Belgium; Department of Biology, Aarhus University, Aarhus, Denmark 8000; Department of Biology, Center for Electromicrobiology, Aarhus University, Aarhus, Denmark 8000

**Keywords:** Microbacterium, extracellular electron transfer, electrode, flavins, oxygen, iron, cable bacteria

## Abstract

*Microbacterium deferre* sp. nov. A1-JK is a metabolically versatile Gram-positive bacterium isolated from the oxic-anoxic interface of freshwater sediment, inhabited by cable bacteria. *M. deferre* A1-JK could simultaneously reduce oxygen and Fe(III), challenging the traditional view of microbial Fe(III)-reduction as a strictly anaerobic process. Electrochemical studies revealed extracellular electron transfer (EET) metabolism facilitated by soluble flavin shuttles, identified via HPLC. Genomic analyses uncovered EET pathways involving cytochrome FccA and flavin reductase FmnA. Its metabolic versatility also allowed for weak electroactivity in alkaline (pH 9–10) and saline conditions (0-4% NaCl). The ability to reduce Fe(III), in the presence of atmospheric oxygen highlights its adaptability to dynamic sediment environments with fluctuating oxygen and Fe(III) gradients. These findings underscore the metabolic versatility of *M. deferre* A1-JK at oxic-anoxic interfaces, providing insights into the coexistence of aerobic and anaerobic processes in a single cell.

**Importance:** *Microbacterium deferre* A1-JK is a newly discovered gram-positive bacterium that employs flavin-mediated extracellular electron transfer to respire minerals and electrodes. Associated with cable bacteria in freshwater sediments, *M. deferre* A1-JK was able to simultaneously reduce oxygen and Fe(III). This makes it highly adaptable to changing environments, such as those found in (cable-bacteria-containing) sediments with fluctuating oxygen and Fe concentrations. The metabolic versatility of *M. deferre* A1-JK offers opportunities to fundamentally revise our understanding of microbial metabolism and nutrient cycling models that strictly separate aerobic and anaerobic metabolisms.

## Introduction

Extracellular electron transfer (EET) is a process in which microbes exchange electrons with extracellular acceptors and donors for metabolic energy generation. As a defining feature of electroactive microbes, EET is often employed to gain access to distant and scarce electron acceptors in the absence of soluble alternatives (*1*). Electroactive microbes can use different mechanisms to perform EET. Direct electron transfer relies on conductive nanowires and outer membrane cytochromes to perform EET via surface contact (*2*, *3*). Mediated electron transfer, on the other hand, uses soluble redox shuttles that diffuse across the cell to ultimately reduce an external electron acceptor (*4–6*).

In recent years, electroactivity has expanded beyond the model organisms *Shewanella oneidensis* and *Geobacter sulfurreducens* (*7*). Several other microbes, which were originally not viewed through the lens of electroactivity have since been discovered to be electroactive (*7*). These include several gram-positive microbes like *Enterococcus faecalis* (*8*, *9*), *Listeria monocytogenes* (*10*), *Lactiplantibacillus plantarum* (*11*) and filamentous *Lysinibacillus varians* GY32 (*12*). Electroactive microbes have been discovered in all kinds of ecosystems (*13*), as electroactivity plays a key role in adapting to nutrient/oxygen limited zones in changing environments (*14*). Sulfidic freshwater and marine sediments populated by cable bacteria are examples of such environments. Cable bacteria are filamentous, sulfide-oxidizing bacteria present ubiquitously in freshwater and marine sediments (*15–17*). Cable bacteria spatially segregate sulfide oxidation from oxygen reduction but link them via long-distance electron transport through conductive periplasmic fibers over centimetre distances (*17*). Distinct biogeochemical gradients of oxygen, sulfide and pH are created in sediments due to the electrochemical activity of cable bacteria, which consume oxygen at a record-breaking rate (*18*). In such sediments, a very dynamic microaerophilic veil is located at the oxic-anoxic interface, with trace amounts of oxygen in the veil. The microaerophilic veil contains (micro)aerobic bacteria, which are often facultative anaerobes (*19*, *20*).

It was discovered that other microbes flock around cable bacteria to indirectly access oxygen via the cable bacteria filaments (*21*). This intriguing phenomenon was proposed to happen through interspecies EET and raises possibilities of studying not only interspecies interactions, but also the role of electroactivity in flocking microbes in such a dynamic environment (*21*, *22*). From the cable-bacteria transversed microaerophilic veil, a novel species *Microbacterium deferre* A1-JK was isolated and studied here for its electroactive metabolism.

*Microbacterium* species are gram-positive and found in diverse environmental conditions including a high-altitude desert environment (*23*), human clinical specimens (*24*) and a radioactive waste site (*25*). As polyextremophiles they can grow under several different conditions, as psychrophiles (*26*), halophiles (*27*), under a wide range of pH conditions (pH 5-11) and as facultatively anaerobic (*28*). In pure cultures, *M. deferre* A1-JK can utilise several carbon sources, as most others of this genus, indicating metabolic flexibility (*29*). Our experiments demonstrate that it can respire anaerobically on Fe(III) and electrodes through flavin-mediated EET. We present the mechanism of EET in *M. deferre* A1-JK along with a potential model for its electron transfer, supported by physiological and genomic analyses. Further, we demonstrate the reduction of Fe(III) by *M. deferre* A1-JK under aerobic conditions, contrary to the widely accepted notion of Fe(III) reduction occurring only anaerobically (*30*). This discovery could have implications for understanding microbial biogeochemical cycling on early Earth transitioning towards the Great Oxidation Event, besides offering opportunities to study bifurcations in microbial metabolism to optimise growth and survival in fluctuating redox-graded environments.

## Results and Discussion

### Current generation by A1-JK in electrochemical reactors

The ability of *M. deferre* A1-JK to use insoluble and soluble electron acceptors was initially evaluated electrochemically. In electrochemical cells, carbon felt electrode constituted the insoluble electron acceptor. All three-electrode cells inoculated with *M. deferre* A1-JK demonstrated a similar trend: a lag phase of around 20 hours as the cells acclimatized to the electrode as the electron acceptor; rising-current phase, leading to a peak; and finally, a decrease in current (Figure 1a). Current levels increased steadily from ∼10 µA at 24 hours and peaked at ∼27 µA after 60 hours. The subsequent decrease in current likely corresponded to a depletion of nutrients, as fresh medium injection into the reactors increased the current to previous levels (Figure 1a). Sterile nutrient medium, heat-killed *M. deferre* A1-JK and *M. deferre* A1-JK without an electron donor did not generate any current, implying that the microbe oxidized the provided electron donor (lactate) and subsequently reduced the electrode.

**Figure 1:**
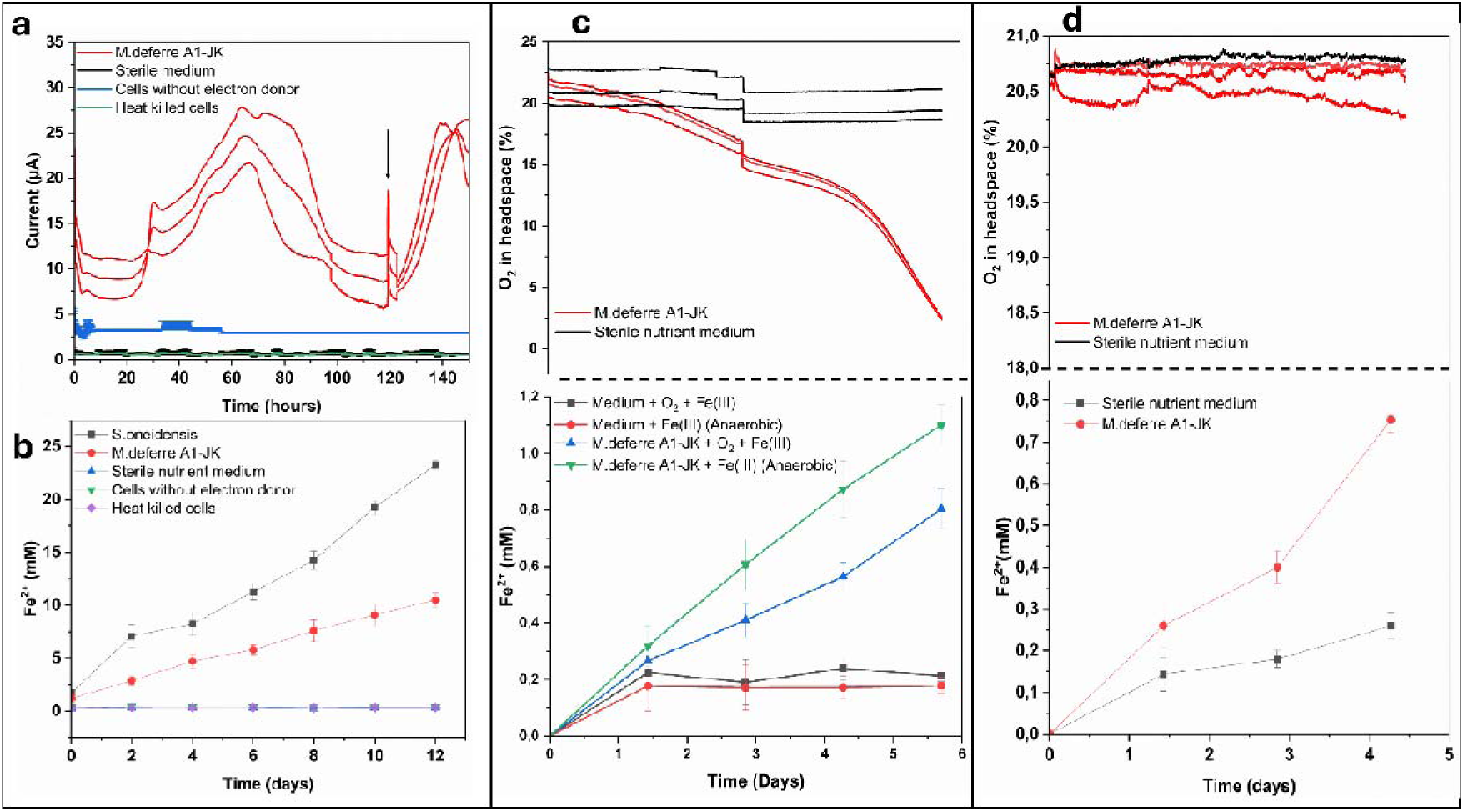
Reduction of insoluble and soluble electron acceptors by *M. deferre* A1-JK. a. Chronoamperometric plots of *M. deferre* A1-JK from the three-electrode cells show a lag phase, log phase with increasing current and a decline in current. The arrow represents the time point at which fresh nutrient solution was injected. b. Reduction of Fe(III) and the subsequent generation of Fe(II), measured by ferrozine assay. c. Simultaneous reduction of oxygen and soluble Fe(III). Oxygen in the headspace was consumed over time (top panel), along with concomitant reduction of Fe(III) (bottom panel). Fe(III) reduction was greater anaerobically. d. Fe(III) reduction under constant oxygen-saturated atmosphere. Top panel: oxygen concentration measurements. Bottom panel: reduction of Fe(III) to Fe(II), measured by ferrozine assay.

### Soluble iron reduction under oxic and anoxic conditions

*M. deferre* A1-JK also demonstrated an ability to reduce soluble Fe(III) to Fe(II) under anoxic conditions, which was determined by the ferrozine assay with ferric citrate provided as the electron acceptor (Figure 1b). Fe(III)-reduction activity was lower than that of *S. oneidensis*, though Fe(III) reduction by *M. deferre* A1-JK increased over time.

The ability to reduce Fe(III) in the presence of oxygen was evaluated since both of these electron acceptors are present in cable bacteria-containing environments from which *M. deferre* A1-JK was isolated. In two parallel experiments, oxygen concentration was measured in the headspace of the culture tubes. In the first set of experiments, after the initial flush of atmospheric air, oxygen concentrations in *M. deferre A1-JK* cultures containing Fe(III)-citrate (Figure 1c, top panel, red lines) exhibited a distinct decline, which initially reflected atmospheric oxygen levels (∼21%). In contrast, oxygen levels in the sterile nutrient medium control (Figure 1c, top panel, black lines) remained stable throughout the experiment. The decrease in oxygen concentration in the *M. deferre A1-JK* cultures indicates active microbial oxygen consumption. In parallel, Fe(III) reduction, measured as Fe(II) accumulation, was assessed under these conditions (Figure 1c, bottom panel, blue line). Fe(II) production was evident in the presence of *M. deferre A1-JK*, with rates dependent on oxygen availability. Under aerobic conditions, Fe(II) accumulation was moderate, indicating that Fe(III) reduction occurred alongside oxygen consumption. In anaerobic conditions, for comparison using the same experimental setup, Fe(II) production was higher than in the presence of oxygen (green line), reaching a maximum concentration of ∼1.2 mM after six days. In sterile medium controls, minimal Fe(II) accumulation was observed under both aerobic and anaerobic conditions.

Figure 1d presents the results of the second set of Fe(III)-reduction experiments conducted under oxic conditions, now with oxygen concentrations maintained constantly at the levels present in atmospheric air (∼21%). Notably, Fe(II) production occurred consistently, even in the presence of oxygen-saturated air. This demonstrates that *M. deferre* A1-JK can simultaneously reduce Fe(III) and utilize oxygen as an electron acceptor, underscoring its metabolic flexibility. These results challenge the long-held notion that Fe(III) reduction is strictly anaerobic and support similar evidence that facultative microorganisms can mediate Fe(III)-reduction in oxic environments (31).

In natural sediment environments, cable bacteria activity leads to creation of microaerophilic zones, with simultaneous precipitation of iron oxides (*32*). In such environments, the ability of *M. deferre* A1-JK to simultaneously couple oxygen and Fe(III) reduction to oxidation of electron donors could be very important for its survival. This phenomenon could involve localized oxygen depletion at the microscale or metabolic strategies such as respiratory chain bifurcation(*33*). This is consistent with findings from *S. oneidensis* (*31*), where microscale gradients allowed Fe(III) reduction under oxic conditions. The prioritization of oxygen as an electron acceptor, likely due to its higher thermodynamic favourability, appears to partially suppress Fe(III) reduction under aerobic conditions (Figure 1c). However, Fe(III) reduction becomes more pronounced when oxygen availability is limited, as observed under anaerobic conditions (Figure 1c, green line). This metabolic flexibility has significant implications for redox-stratified environments, such as sediments, aquifers, and soils, where oxygen and Fe(III) gradients coexist. By simultaneously reducing oxygen and Fe(III), *M. deferre* A1-*JK* could potentially contribute to the generation of a specific niche for iron cycling and thus influence geochemical processes, the community composition and distribution in these habitats. Additionally, the resilience of *M. deferre* A1-JK to oxygen fluctuations makes it a promising candidate for applications in bioremediation of metal-contaminated environments and bioelectrochemical systems.

### Mechanism of extracellular electron transfer

Cyclic Voltammetry (CV) of *M. deferre* A1-JK cultures in the presence of a carbon felt working electrode produced a distinct redox couple at −0.302 V (Figure 2a), suggesting the involvement of soluble redox shuttles mediating electron transfer (*5*). The CV shows a second oxidation peak, occurring at +0.18 V (Figure 2a). This peak was, however, inconsistent in Differential Pulse Voltammetry (DPV) (Figure 2b), suggesting a smaller role in EET. Based on the potential, this peak was identified as soluble iron from the nutrient medium as demonstrated previously (*34*).

**Figure 2.**
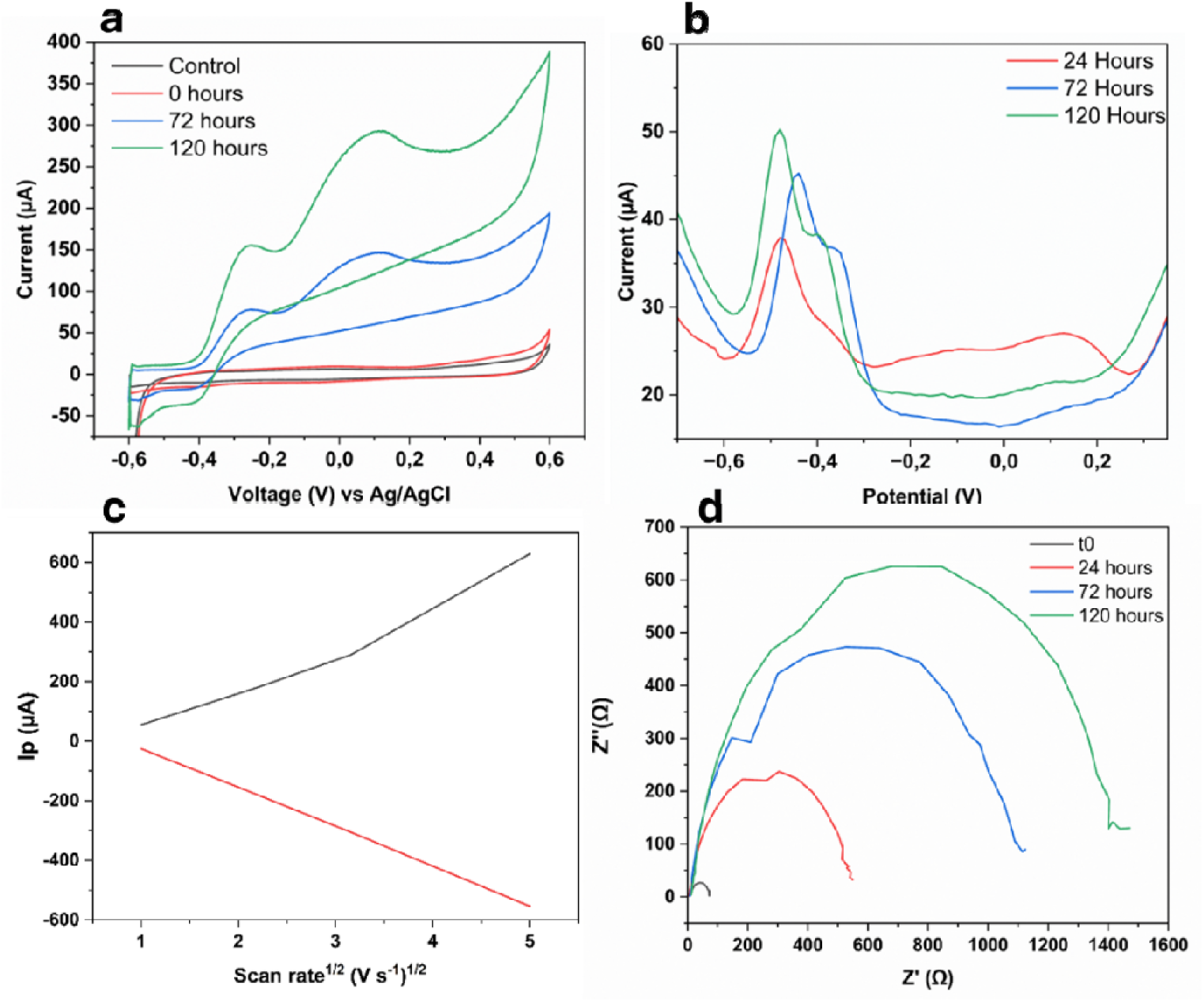
a. Cyclic voltammograms of *M. deferre* A1-JK from electrochemical cells revealed redox couples corresponding to EET mediators at −0.302V. b. Differential pulse voltammetry produced peaks that increased over time, suggesting the involvement of redox shuttles. c. Randles–Ševčík plot of *M. deferre* A1-JK cultures confirmed that electron transfer occurred via diffusion. d. EIS plots for *M. deferre* A1-JK at different time points during their growth in the three-electrode cell.

The peak heights at negative potentials increased in the CV and DPV over time, indicating a rise in redox shuttle concentrations (Figure 2b). This suggested that there was a synthesis of potential redox shuttles, with concentrations correlated to cell growth. Randles–Ševčík plots (Figure 2c) based on differing scan rates produced linear curves proportional to the square root of the scan rates, confirming that electron transfer occurred by diffusion and that mediator concentration was the limiting factor in current generation.

Electrochemical Impedance Spectroscopy (EIS) analysis corroborated the findings on diffusion-based electron transfer (Figure 2d). As the cells in the reactor acclimatized to reduce the electrode, the resistance to charge transfer decreased, with the smallest resistance corresponding to the highest current generation. Interestingly, the development of biomass on the anode over time did not reduce charge transfer. This suggests that there may not have been a strong biofilm formation, and that the cells depended on mediators to perform electron transfer. Biofilm assays and SEM imaging on electrodes revealed weak ability of *M. deferre* A1-JK cells to form biofilms (Figure S1, S2).

### Media swap and HPLC confirm flavin-mediated EET

By separating the electrolyte (spent medium without the cells) and the carbon-felt electrodes from the three-electrode cells we determined the degree of direct and mediated electron transfer by *M. deferre* A1-JK. A schematic is provided in Figure S3 to clarify the media swap experimental setup. DPV on reactors with fresh medium and used electrodes containing attached cells of *M. deferre* A1-JK showed no redox peaks, implying an absence of components necessary for performing direct electron transfer (Figure 3a). Redox peaks were clearly present with the *M. deferre* A1-JK media previously used in a three-electrode cell, confirming the presence of secreted mediators.

**Figure 3.**
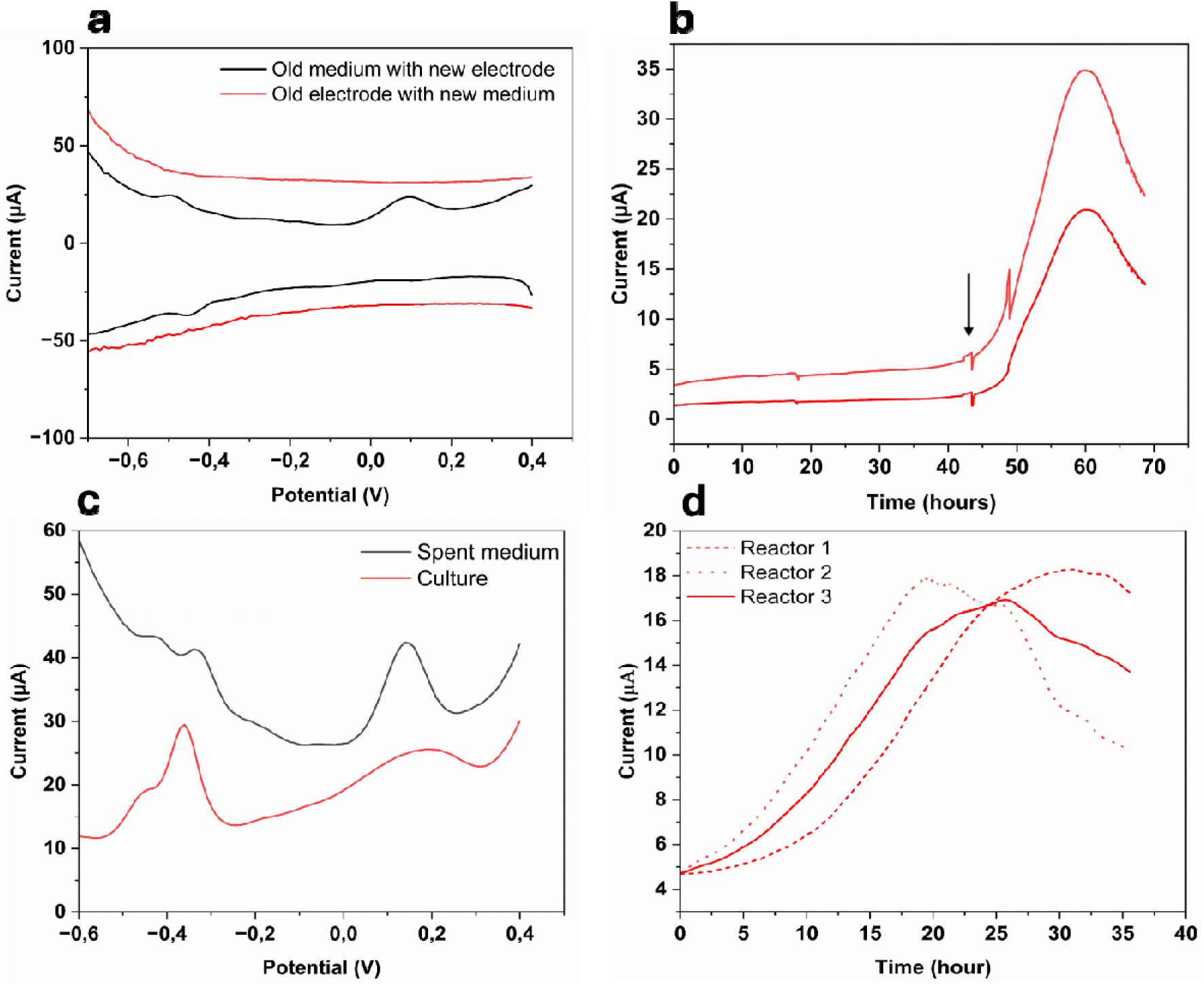
a: DPV of the *M. deferre* A1-JK old electrode with a new medium (red) and *M. deferre* A1-JK nutrient medium with a new electrode (black) revealed redox-active peaks in the medium, indicating the presence of secreted redox shuttles. Mean values from triplicates are presented. b. Current instantly increased during chronoamperometry with external addition of 2 µM riboflavin (arrow) in reactors with used *M. deferre* A1-JK electrodes in fresh nutrient medium. Duplicates are presented as the third reactor was damaged. c. DPV of spent medium and *M. deferre* A1-JK culture from the three-electrode cell reveal peaks at similar potentials, confirming mediated EET. Mean values from triplicates are presented. d. *M. deferre* A1-JK inoculated in spent media supplemented with lactate showed a rapid increase in current without any lag time by utilizing flavins already present in the medium.

HPLC analyses of the cell-free spent medium confirmed the presence of riboflavin. This is consistent with reports of flavins occurring at similar negative potential in other studies (*5*, *35*, *36*). Riboflavin concentrations in *M. deferre* A1-JK samples increased from 200 nM after 24 hours to 1.6 µM after 6 days at peak current generation (Figure S4). A test run of *S. oneidensis* MR-1 in the same electrochemical setup yielded riboflavin at a concentration of 2.2 µM. Further, in reactors containing fresh electrodes in used electrolytes, riboflavin concentration was estimated at 1.4 µM. In reactors with fresh nutrient medium and previously used electrodes containing attached cells of *M. deferre* A1-JK, riboflavin concentration after three days was only 100 nM, indicating that redox shuttles were secreted in the medium and direct electron transfer was minimal. When riboflavin (2 µM) was injected into reactors containing previously used electrodes in fresh media, current generation increased from 5 µA to 35 µA (Figure 3b), confirming mediated electron transfer as the mode of EET.

Further, following the end of the experiment, the electrolyte was carefully collected and centrifuged. The supernatant was filtered, and the resulting spent media was analysed via DPV (Figure 3c). Similar peaks were seen in the culture as well as the spent medium, validating the above results. Filter-sterilized spent media supplemented with 20mM lactate was then inoculated with *M. deferre* A1-JK cells in new electrochemical reactors with fresh electrodes (Figure 3d). Current increased rapidly with no lag phase and reached a peak value around ∼20 hours, as the spent media already had sufficient flavins. This contrasts nicely with Figure 1a, where a longer lag phase was observed due to the absence of flavin build-up in a fresh medium. These results further confirm riboflavin-mediated EET. In addition, flavins were secreted even under oxic conditions, with riboflavin concentrations of 0.5 µM secreted under aerobic growth conditions. Riboflavin secretion in the presence of oxygen could explain reduction of Fe(III) in oxic conditions, as described earlier.

### Versatility to perform EET in alkaline and halophilic conditions

Because *M. deferre* A1-JK could encounter varying pH conditions in its natural environment, its capability to grow at a wide pH range between pH 5 and pH 10 was tested. In the oxic zone, where iron oxides are formed, the pH can be strongly alkaline (up to pH 9.0) due to cable bacteria activity (*37*). Chronoamperometric plots at pH 9 and 10 for *M. deferre* A1-JK cultures showed increasing currents, indicating weak electroactivity (Figure S5). To determine whether *M. deferre* A1-JK was able to grow at different pH range (5–10), growth curves under aerobic conditions were done (Figure S6). These showed that *M. deferre* A1-JK was adaptable to pH as long as oxygen was present.

*M. deferre* A1-JK also demonstrated weak electroactivity under 4% salt conditions (Figure S5). Specific reasons behind the functionality of performing EET under high salt concentrations, remain unknown. *M. deferre* A1-JK was able to grow at 4% NaCl concentrations but did not grow at 10% NaCl (table S1). The closest phylogenetic relative to M. *deferre* A1-JK, *Microbacterium sp*. Gd 4-13, demonstrated the ability to grow at salt concentrations of 10% NaCl (Figure S7) (*27*).

### Potential EET pathways

Potential EET pathways of *M. deferre* A1-JK based on genomic data are highlighted in Figure 4. Its flavin production was reflected in the genome by genes for the flavin synthesis cascade: RibBA,D,E1,E2 (Table S2). A partial route using the periplasmic proteins DmkA,B, FmnA and Ndh2 might control electron fluxes towards aerobic respiration and EET (*38*) as previously seen for other gram-positive electroactive microbes like *L. monocytogenes, L. lactis, L. plantarum, E. faecalis,* and other Gram-positive microbes with FLEET (flavin-based EET) locus (*10*, *39*, *40*). However, in contrast to other FLEET-based electroactivity, the genes for PplA, FmnB and EetA,B, were absent in *M. deferre* A1-JK. Other genes encoding proteins significant to EET, including those associated with the MenBCE complex and FccA were detected in the genome (*40*). The cytochrome FccA would be used to direct electron flow from the quinone pool toward the reduction of riboflavin and FAD via FmnA within the cytoplasm. This is supported by predictions indicating that the FccA proteins are localized to the cytoplasm and the cytoplasmic membrane (Table S2). In previous studies where FccA and FmnA were genetically knocked out in *S. oneidensis*, Fe(III) and fumarate reduction were suppressed, showing a clear involvement of FccA in EET (*41*). *M. deferre* A1-JK was also able to grow with fumarate as the sole electron acceptor under anaerobic conditions (data not shown).

**Figure 4:**
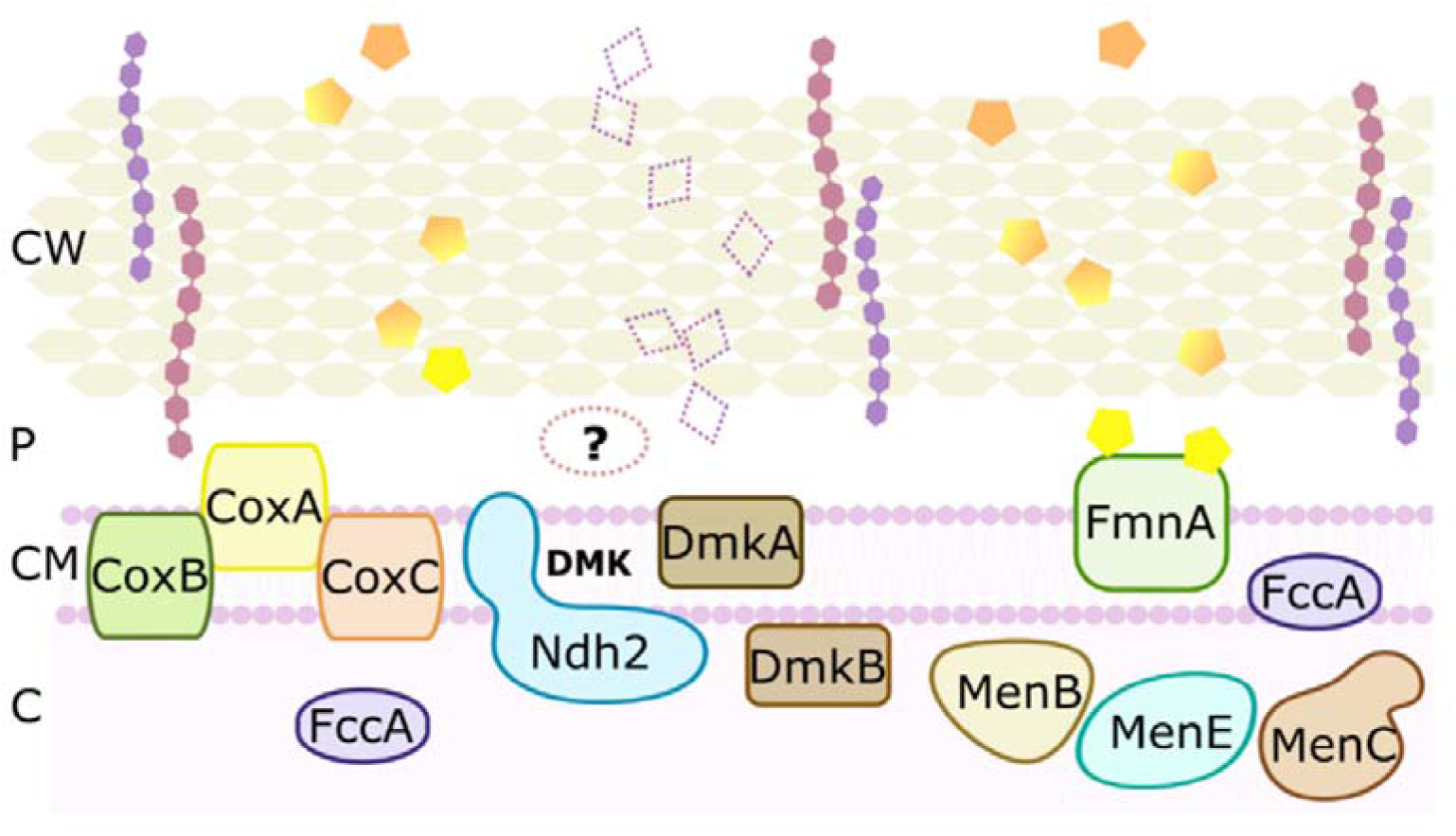
Potential extracellular electron transfer pathways in the genome of *M. deferre* A1-JK. DmkA, DmkB and Ndh2 regulate electron flux towards EET. FccA likely directs electrons towards reduction of riboflavin and FAD via FmnA. Reduced flavins (yellow pentagons) diffuse across the cell wall towards extracellular acceptors. Phenazines (empty diamonds) were not observed experimentally, though genes for phenazine synthesis were observed. CW: cell wall, P: periplasm, CM: cytoplasmic membrane, C: cytoplasm, hexagonal strings represent teichoic acids

The synthesized flavins would be externalized via diffusion through the cell wall, as experimentally observed in the Randles–Ševčík plots (Figure 3c). The flavin exporter YeeO, was encoded in the genome and localised to the cytoplasmic membrane (Table S2), though it has not been reported to be used in electron transport over inner cell membranes or cell walls (*42*). However, recordings of flavin transporters in Gram-positives are becoming more common, and knock-out studies might further elucidate their specific role in metabolic pathways (*22*, *43*).

Reduced flavins are exported to transfer electrons towards extracellular electron acceptors. Outside of flavins, a partial synthesis pathway for phenazines, PhzD,E,G was detected (Table S2). Potentially, both flavins and phenazines could be used as mediators in EET. Yet, only flavin signature peaks were detected in voltammetry measurements. The complex production and energetically expensive costs of phenazines likely prevented their synthesis in the low-energy nutrient medium that was used for the electrochemical cells, or the absence of PhzA,B,F may prevent phenazine synthesis. Another reason for the lack of phenazines could be that some of the genes from the phenazine synthesis operon were used in alternative biosynthetic pathways (*44*).

Interestingly, the outer membrane cytochrome ExtH, discovered in *G. sulfurreducens*, was encountered twice in the genome of *M. deferre* A1-JK (table S2). The finding of a Gram-negative outer membrane protein that forms an EET-related protein complex in a Gram-positive bacterium is notable. In *G. sulfurreducens*, ExtH is involved in selenium uptake (*45*, *46*). ExtH-like proteins in *M. deferre* A1-JK are predicted to be localised to the cytoplasm (Table S2), and expectations are that these cytochrome homologs support the electron transfer to Ndh2 and to DmkA and DmkB. However, the specific role in EET of these cytochromes in *M. deferre* A1-JK remains unclear. We propose the existence of a periplasmic protein that helps to facilitate fast internal electron transport, although no clear evidence has been identified. This hypothesis could explain the absence of the signature peak in DPV. Further studies that involve knock-out mutations are required to elucidate the role of the ExtH-homologs, the presence of a periplasmic transport protein, and any other associated proteins.

### Metabolic potential of M. deferre A1-JK

As stated above, *M. deferre* A1-JK has several potential pathways to interact with extracellular electron acceptors that were discovered in the 3.5 Mb-large genome, consisting of 5 contigs with a coverage of 334, and GC% of 69.5. With the large metabolic versatility documented in the *Microbacterium* genus we investigated the genomic potential of *M. deferre* A1-JK towards drug degradation. Genomic analyses revealed the ability to degrade several different types of drugs, polystyrene, DMSO_2_ (dimethylsulfone or methylsulfonylmethane; MSM) and formaldehyde (Figure S8). Further, the capability of breaking down (poly)sugars which it likely uses for assimilation, was also present, though growth on L-oriented amino acids is likely not possible. Assimilation of single or dual-chain sugars occurs only after 48-72 h, while fermentation of such sugars could occur after 24 h. *M. deferre* A1-JK was not capable of producing H_2_S but may produce very small amounts of N_2_ (Table S1).

The ability to degrade various drugs could have implications for environmental and biotechnological applications. The ability to break down complex organic compounds highlights *M. deferre’*s potential role in bioremediation, particularly in the degradation of persistent pollutants. Additionally, its capacity to assimilate and ferment (poly)sugars, may support its use in waste treatment processes where diverse organic substrates are present. Further exploration of its metabolic pathways could advance our understanding of its ecological niche and its utility in industrial processes.

## Conclusions

*M. deferre* A1-JK is the first reported Gram-positive electroactive bacterium within the *Microbacterium* genus. It has a versatile metabolism and it can use many different carbon sources, including (poly)sugars. A flavin-mediated EET strategy is employed by *M. deferre* A1-JK under both aerobic and anaerobic conditions. The ability to perform mediated EET is present even in the presence of atmospheric levels of oxygen and harsh conditions, like high pH 9-10 and 4% salt. However, it appears to have an incomplete set of EET proteins, compared to the known FLEET genes or other Gram-positive EET genes. This leaves the possibility of novel proteins in *M. deferre* A1-JK, that are used to support the electron transport.

In sediments where cable bacteria are found, unique selection pressures like pH and oxygen fluctuations arise because of cable bacterial activity. Such an environment could create the necessary conditions for *M. deferre* A1-JK to thrive by performing EET towards minerals available as electron acceptors, as well as reducing oxygen to optimise its growth rate. The demonstration of EET in the presence of oxygen warrants further studies on metabolic capabilities and adaptations in response to changing environmental conditions. To better understand the molecular mechanisms enabling simultaneous oxygen and Fe(III) reduction, studies using gene knockouts and transcriptomics are critical. Metabolic modelling could elucidate the energy trade-offs and regulatory pathways involved. Such studies will also be useful in further understanding its influence on the biogeochemistry and its role in EET-related nutrient cycling. It may be worth examining the cell wall and associated proteins for any unique structural adaptations to combat high alkalinity and salt concentrations. These insights would advance ecological understanding, the design of engineered systems for sustainable biotechnological applications and fundamentals of microbial bioenergetics.

### Description of M. deferre A1-JK sp. nov

*Microbacterium deferre* (Lat. from *deferre/defero*; reduce, transport or exchange, after its capabilities to transport and exchange electrons through the reduction of electron shuttles). Cells are gram-positive, irregular and non-motile rods that become ovoid shaped when older (1.1-1.8 µm × 0.4-0.7 µm). They form mucoid, faded-yellow colonies (3-6 mm diameter) but colony color and size differ between growth media. *M. deferre* A1-JK is catalase positive, and OF-test positive for glucose, arabinose and sucrose and weakly so for mannose, maltose and gluconate. It is capable of growing on a wide range of complex growth media and may produce low concentrations of N_2_. NaCl concentrations of 10.0% (w/v) are very poorly tolerated, with normal growth occurring until at least 4.0% NaCl. Growth occurs aerobically or anaerobically at a pH range from 5.0-10.0, and at a temperature range from 4°C to 37.5°C. The size of the genomic DNA is ∼3.5 Mb, with a G+C content of 69.32%.

*M. deferre* A1-JK was isolated from a microaerophilic veil that formed in a trench slide containing freshwater sediment with a single strain of cable bacteria: *Electronema aureum* GS(*47*, *48*). The sample was extracted with a glass capillary in March 2019 and cultivated until a pure culture was reached in June 2019. The accession number for *M. deferre* A1-JK genome sequence is PRJNA1125322.

## Materials and Methods

### Isolation

A trench slide was used to inoculate sediment containing freshwater cable bacterium *Electronema aureum* GS (*48*) as per Bjerg *et al* (*17*). Microaerophilic veils are characteristically formed in trench slides as oxygen diffuses into the trench slide. After cable bacteria were observed stretching out from the trench, visualized capillary suction was employed to draw out the liquid from the microaerophilic veil, as described earlier by Lustermans *et al* (*49*). Following plating on Reasoners 2-A agar plates, *M. deferre* A1-JK colonies were identified and transferred under laminar air flow to sterile agar plates to obtain pure cultures. The agar plates were incubated aerobically in dark conditions at 15 °C.

### Physiological testing

API 20E and 20NE test strips (Biomérieux) were inoculated and read according to the manual. Strips were incubated in the dark at room temperature for 24 hours initially, and subsequently for 48 and 72 hours. Growth was determined at different temperatures: 4°C, 15°C, 18-22°C (room temperature), and 37.5°C, by inoculating and incubating Nutrient Broth (NB), VWR, R2A (Millipore) and Luria-Bertani (VWR) agar plates in the dark. Similarly, growth in incremental salt concentrations 1-4%, and 10% NaCl in LB plates were tested. Since LB contains 1% NaCl, R2A plates were used for 0% NaCl.

### Nutrient medium composition for electrochemical cells

A defined nutrient medium was prepared for electrochemical studies. The nutrient medium was modified *Shewanella* basal medium (SBM), consisting of the following components (per litre): K_2_HPO_4_, 0.225 g; KH_2_PO_4_, 0.225 g; NaCl, 0.46 g; (NH_4_)_2_SO_4_, 0.225 g; MgSO_4_·7 H_2_O, 0.117 g; HEPES, 23.83 g. Sodium lactate (20 mM) was provided as the carbon source. Mineral and vitamin mix (5 ml each) was added. The mineral mix consisted of nitrilotriacetic acid, 1.5 g; MnCl ·4 H_2_O, 0.1 g; FeSO_4_·7 H_2_O, 0.3 g; CoCl_2_·6 H_2_O, 0.17 g; ZnCl_2_, 0.1g; CuSO_4_·5 H_2_O, 0.04 g; AlK(SO_4_)_2_ ·12 H_2_O, 0.005 g; H_3_BO_3_, 0.005 g; Na_2_MoO_4_, 0.09 g; NiCl_2_, 0.12 g; NaW_4_ ·2 H_2_O, 0.02 g and Na_2_SeO_4_, 0.1 g (per litre), while the vitamin mix consisted of folic acid, 0.002 g; biotin, 0.002 g; pyridoxine HCl, 0.02 g; thiamine, 0.005 g; nicotinic acid, 0.005 g; pantothenic acid, 0.005 g; cyanocobalamin, 0.0001 g; p-aminobenzoic acid, 0.005 g and thioctic acid, 0.005 g.

### Iron reduction assay (Ferrozine assay)

Fe(III) reduction was estimated using ferrozine assay as described earlier(*50*). To summarize, *M. deferre* A1-JK was grown overnight in LB medium and washed and resuspended in the amended SBM to an OD of 1. Thirty microliters of this suspension was added to 270 µl of SBM containing vitamins, minerals, 30 mM lactate and 30 mM Fe(III) citrate in 96-well plates. The plates were flushed with nitrogen for 20 minutes. 50 µl of 5M HCl was added to prevent oxidation of Fe(II). 30 µl was taken from this well and added to 270 µl of 0.5M HCl. 50 µl of this diluted sample was mixed with 300 µl of ferrozine reagent. Absorbance was measured at 562nm. Standard curves were made from FeSO_4_ dissolved in 0.5M HCl. Fe(III) reduction was compared with *S. oneidenses* MR-1 grown in SBM with 30 mM lactate as the electron donor.

### Electrochemical cell construction

Three-electrode cells fabricated from glass bottles (30ml working volume) were used as electrochemical reactors. The working electrode consisted of carbon felt (Thermo Fisher, dimensions 2 cm × 2 cm × 0.3 cm), Ag/AgCl (REF201 Radiometer Analytical, Denmark) constituted the reference electrode and a Pt wire was used as the counter electrode. The cells were inoculated with 3 ml (10% inoculum) of *Microbacterium* culture (OD 0.4), followed by addition of 27ml nutrient medium. After inoculation, nitrogen was sparged for 20 minutes to make the reactors anoxic, following which electrochemical experiments were conducted. The reactors were operated at 25 °C.

### Electrochemical measurements

A potential of 200 mV was applied using a MultiSens4 (PalmSens) potentiostat for chronoamperometry (CA). Additionally, chronoamperometric measurements were performed under alkaliphilic and halophilic conditions (pH 8-10; 4% salt). Cyclic voltammetry (CV) was performed at a potential window from −0.6 V to +0.6 V at a scan rate of 1 mV. For Randles-Sevcik plots, CV was done at varying scan rates of 1 mV, 5 mV, 10 mV and 25 mV. Differential pulse voltammetry (DPV) was performed with the same scan window as cyclic voltammetry, using 1 s period, 0.3 s pulse time, 50 mV pulse size and 1 mV step size. EIS was done in potentiostatic mode based on midpoint potential of the cyclic voltammogram (−0.384 V). The potential amplitude was 30 mV.

At the end of the experiments, spent medium was analysed via CV and DPV. The electrolyte from the three-electrode cells was collected and centrifuged at 6000 g for 15 min, followed by collecting the supernatant and repeating the centrifugation for 30 min. The resulting supernatant was filtered using a 0.22 µm membrane to obtain cell-free spent medium. The spent medium was transferred to new three-electrode cells with fresh electrodes for electrochemical studies.

CA, CV, DPV, Randles-Sevcik plots and EIS were all conducted in triplicates.

### Media swap experiments

After operating the three-electrode cells for a period of 7 days, the electrodes and electrolyte were separated in fresh reactors to ascertain redox-active components to elucidate the EET mechanism. The electrodes were carefully transferred to new electrochemical reactors with fresh nutrient medium, while fresh electrodes were introduced into the old electrolyte.

### Aerobic Fe(III) reduction experiments

*M.deferre* A1-JK was inoculated in the above-described nutrient medium in Hungate tubes (final volume 10 ml) with lactate as the electron donor. Fe(III) citrate (2 mM) was introduced in the medium as an electron acceptor. The starting O.D. of the culture after inoculation was 0.4. The culture and headspace in the Hungate tubes were sparged with air for 20 minutes to make it oxygen saturated. The headspace of the Hungate had optical oxygen spot sensors that were connected to a Firesting O_2_ device (Pyroscience). Data was logged using Pyro Oxygen Logger software. O_2_ concentrations were continuously measured in the headspace by oxygen spot sensors, while Fe(III) reduction was estimated by the ferrozine assay as described earlier. In a separate experiment, air was continuously sparged using an aquarium pump to keep oxygen levels constant at atmospheric levels. Fe(III) reduction was estimated by ferrozine assay as described earlier.

### HPLC

HPLC was done to quantify redox shuttles. Based on the redox peaks in cyclic voltammograms, flavins were targeted for quantification by HPLC. Filtered spent medium from three-electrode cells was used. HPLC was done on a Dionex UltiMate 3000 UPLC system (Thermo Scientific) equipped with a C18 analytical column. Fluorescence of riboflavin was monitored (excitation 440 nm; emission 525 nm). Final riboflavin concentration was determined by eluting a series of riboflavin standards (100 nM, 200 nM, 400 nM, 800 nM, 1 µM, 2 µM and 3µM) and preparing a standard curve. Samples were run in triplicates.

### Microtiter Dish Biofilm Formation Assay

The ability of *M. deferre* A1-JK to form biofilms was estimated by the microtiter dish biofilm formation assay, as described previously (*51*, *52*). Briefly, the cells from the three-electrode reactors were diluted 1:100. 100µl of this culture was added in 96-well plates, followed by addition of 125 µl of 0.1% crystal violet solution. Following incubation at room temperature for 15 minutes, the plates were washed with distilled water. After drying, 125µl of 30% acetic acid was added in the wells. Absorbance was measured at 550nm.

### Gram staining

Cells of *M. deferre* A1-JK were deposited onto a drop of milliQ water, airdried and heat-fixated on a microscopy slide by moving through a flame. The sample was stained with crystal violet for 1 min and thoroughly rinsed with milliQ water, followed by iodine solution flooding for 1 min and rinsing. Decolouring was done for 2 s with 100% ethanol and rinsing. Safranine solution was used for counterstaining for 30 s and rinsing. The slide was airdried and inspected with a light microscope as described below.

### Light and Scanning Electron Microscopy (SEM)

Plate culture of *M. deferre* A1-JK was put on glass microscopy slides, anoxic tap water was added and covered with a glass coverslip. Light microscopy images were made with phase-contrast at 1000x with an oil lens (Zeiss Axioplan 2), black and white camera (Exi Blue). SEM samples were prepared by putting a carbon sticker on an aluminium SEM pin in a dust free environment. Carbon felt electrodes (after concluding the electrochemical cell measurements) were deposited on the carbon sticker and dried for 24 h, followed by vacuum gold-sputtering for 40s (Agar Sputter Coater). Images were made with a Phenom ProX SEM (Phenom-World BV) using a backscattered electron detector with a 10 kV electron beam. Images were used to determine motility, spore formation, and measure cell size. Sizes were measured using the measure tool in Fiji’s ImageJ (*53*).

### DNA extraction, 16S gene- and genome sequencing

As described in Lustermans *et al*(*49*), 16S rRNA-based taxonomy was determined through a colony PCR on a single colony of *M. deferre* A1-JK which was Sanger sequenced (Macrogen). A second sample of multiple colonies of *M. deferre* A1-JK (also from an agar plate) was used for DNA extraction with the DNeasy PowerSoil kit (Qiagen). Concentration and length quality checks were done using Qubit (Fischer Scientific) and an Implen Spectrophotometer (Fischer Scientific) according to the manual. Genomic DNA was sent for Illumina MiSeq sequencing 2×150bp (Eurofins, Germany).

### Genome assembly, annotation, analyses and phylogeny

Trimming, size filtering and adapter removal was done with Trimmomatic v0.39 crop 125 and headcrop 20 (*54*). Trimmed paired-end reads were *de novo* assembled with SPAdes v3.15.5 with parameters(*55*) isolate, default coverage cut-off and default k-mer values. Statistics for completeness, contamination, quality, G+C content and coverage were calculated with Quast v5.2.0 (*56*), fastq-stats v1.01 and CheckM2 v1.0.1(*57*),(*58*).

All available *Microbacterium* genomes were downloaded from NCBI genbank and used as database for gene identification with PROKKA v1.14.5 (*59*). Before use, the database was deduplicated with CD-HIT v4.8.1 with a length cut-off of 0.8, clustered into the most similar cluster, and 0.9 sequence identity threshold (*60*, *61*).

Metabolic pathways were determined with BlastKOALA and KEGG mapper (*62*, *63*). EET- and electron shuttle synthesis genes were identified through blastp (*64*), using a manually, in-house curated, protein database of EET-related genes with an E-value cut-off of 1E-6. Cellular location predictions were done on the amino acid sequences with positive blastp results with psortb v3.0.3 (*65*).

Phylogeny and taxonomic trees were determined with GTDB-Tk 2 infer and align (*66*), FastANI (*67*), and Figtree2 (http://tree.bio.ed.ac.uk/software/figtree/) on all *Microbacterium* genomes from NCBI, or a subset of the phylogenetic tree: a branch which contained A1. As outgroup, *Mesorhizobium loti* LMG 6125 was used. ANI (http://enve-omics.ce.gatech.edu/ani/index) was used to determine differences with the two closest relatives *M. hatanonis* JCM 14558 and *M. sp.* Gd 4-13.

The genome of *M. deferre* A1-JK is accessible at NCBI genbank under the reference PRJNA1125322.

## Acknowledgments

KA and JJML both acknowledge funding from the European Union’s Horizon Research and Innovation Program under the Marie Sklodowska-Curie grant agreement (project 101109777; KA and 101152850; JJML). The authors thank Dimitri Stamatis for his assistance with the HPLC, Lars Børregård Pedersen for assistance with oxygen monitoring experiments and Jeanine Geelhoed for logistical sequencing support. The Danish National Research Foundation (Center for Electromicrobiology, DNRF136) is also acknowledged.

## Author Contributions

Conceptualization: KA, JJML; Methodology and Investigation: KA, NB, LD, JJML; Writing-Original Draft: KA and JJML; Writing-Review & Editing: KA, NB, LD and JJML; Funding Acquisition: KA and JJML

