## Supplementary Information for "Flavin-based extracellular electron transfer under oxic conditions by Gram-positive *Microbacterium deferre* sp. nov. A1-JK in redox-graded environments"

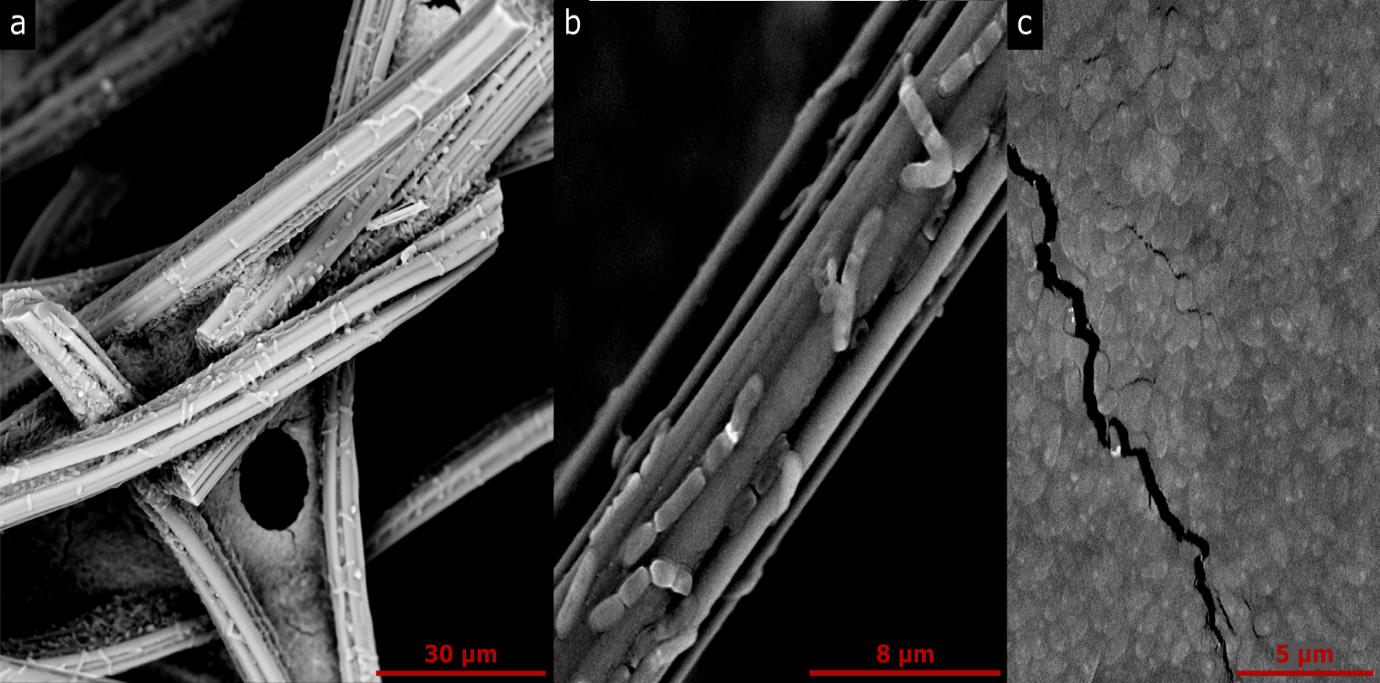

Figure S1: Scanning electron microscopy images of *M. deferre* A1-JK depicting elongated ovoid-rod cells on carbon felt electrodes (a, b). Cell morphology and sizes of *M. deferre* A1-JK cultures on carbon felt electrodes were compared through SEM. Cells were seen as elongated ovoid-rods often as strings of 2-4 cells, and cell size recorded was 1.799 ± 0.35 × 0.672 ± 0.06 µM. Cells aggregated on the electrode, though a thick biofilm was not observed.

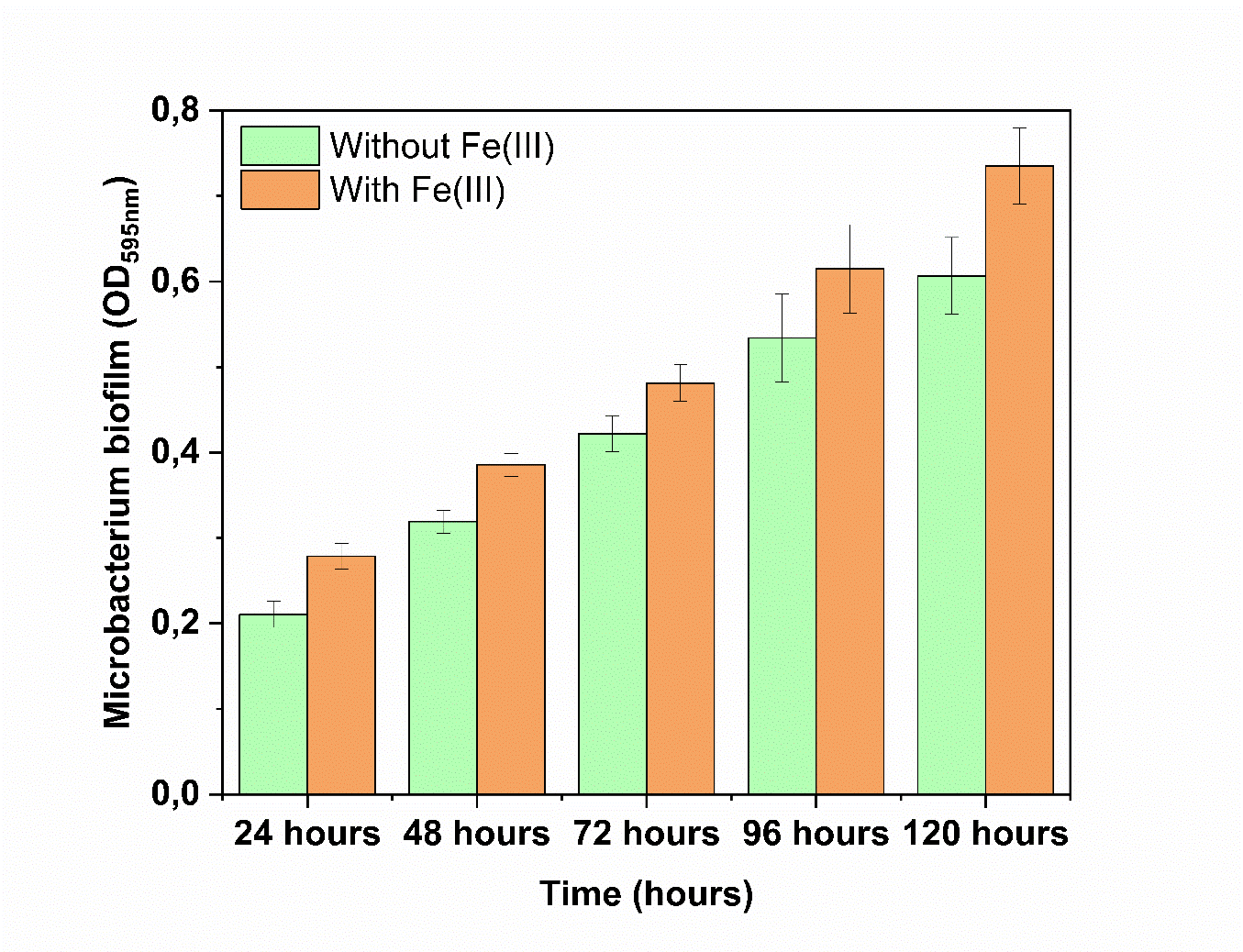

Figure S2: Biofilm formation in *M. deferre A1-JK*, estimated by the microtiter dish biofilm formation assay revealed weak biofilms. Biofilm formation increased in the presence of Fe(III).

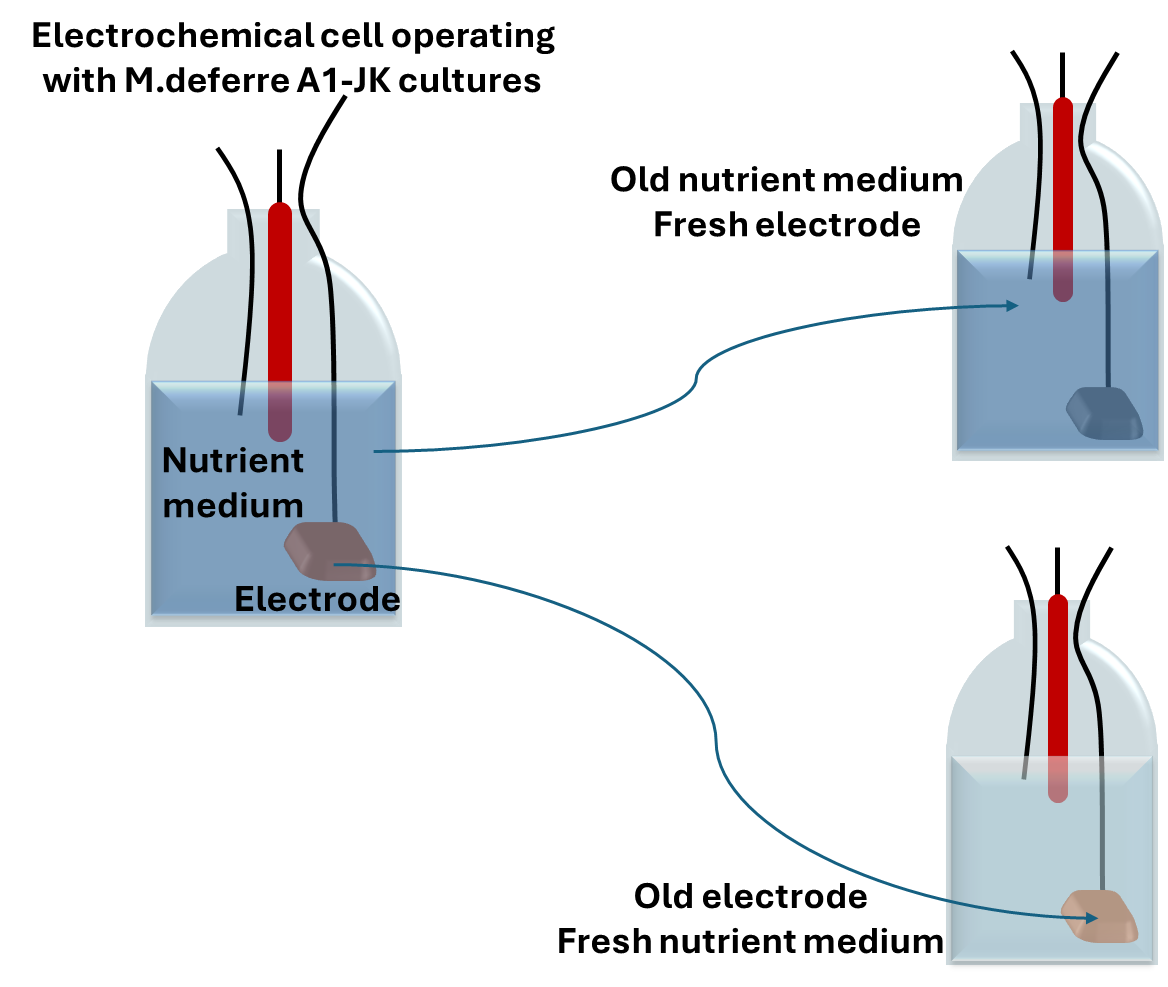

Figure S3: Schematic representation of media swap experiments. The electrode and nutrient medium from *M.deferre* A1-JK electrochemical cells were separated into different reactors to evaluate the role of direct and mediated EET.

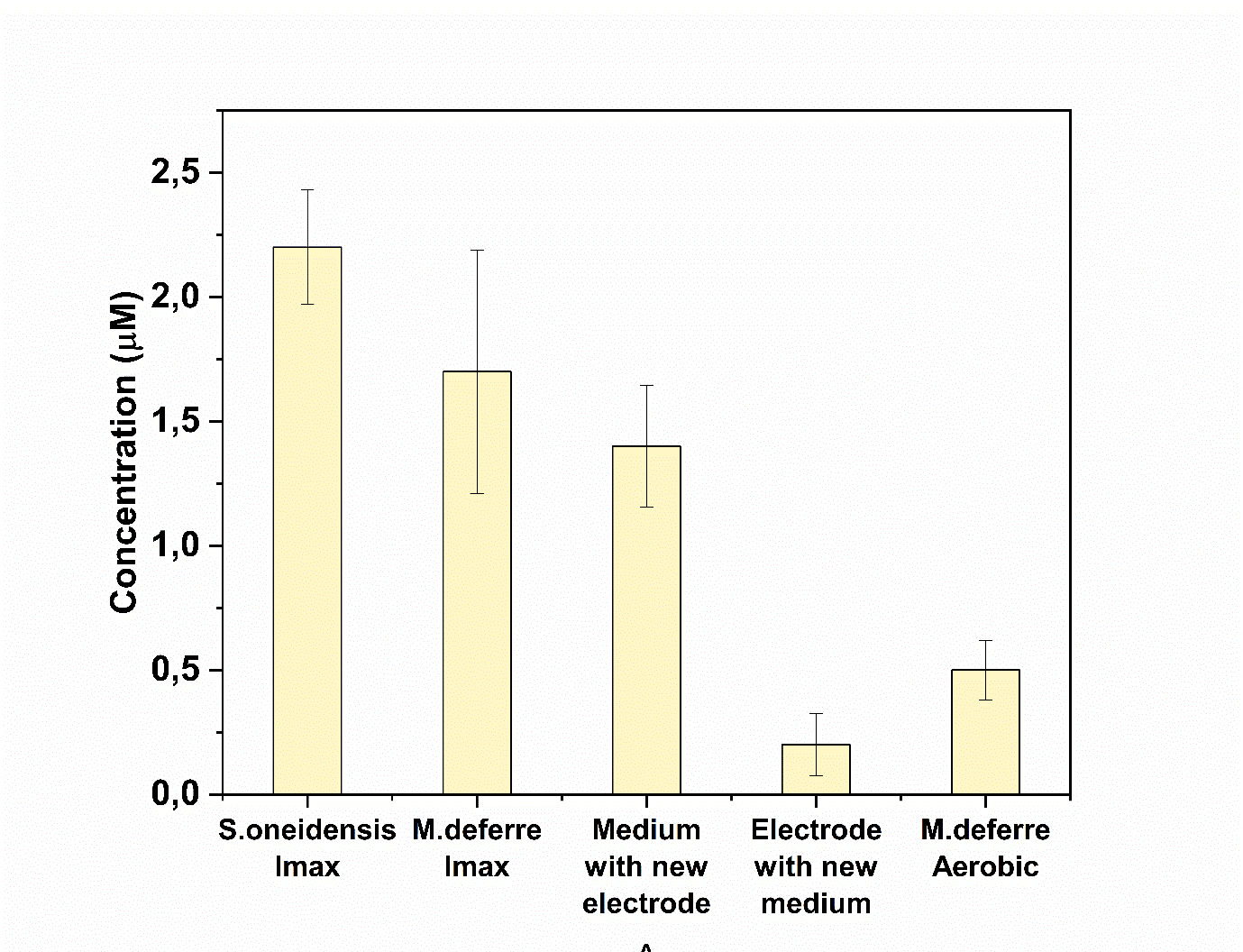

Figure S4: Riboflavin quantification via HPLC. At peak current (Imax), *M. deferre* A1-JK produced 1.6 µM of riboflavin, compared to 2.2 µM produced by *Shewanella* *oneidensis* MR-1. In the media swap experiments, riboflavin was present in the previously used media, indicating mediated electron transfer. Under aerobic conditions, *M.deferre* A1-JK secreted 0.5µM of riboflavin.

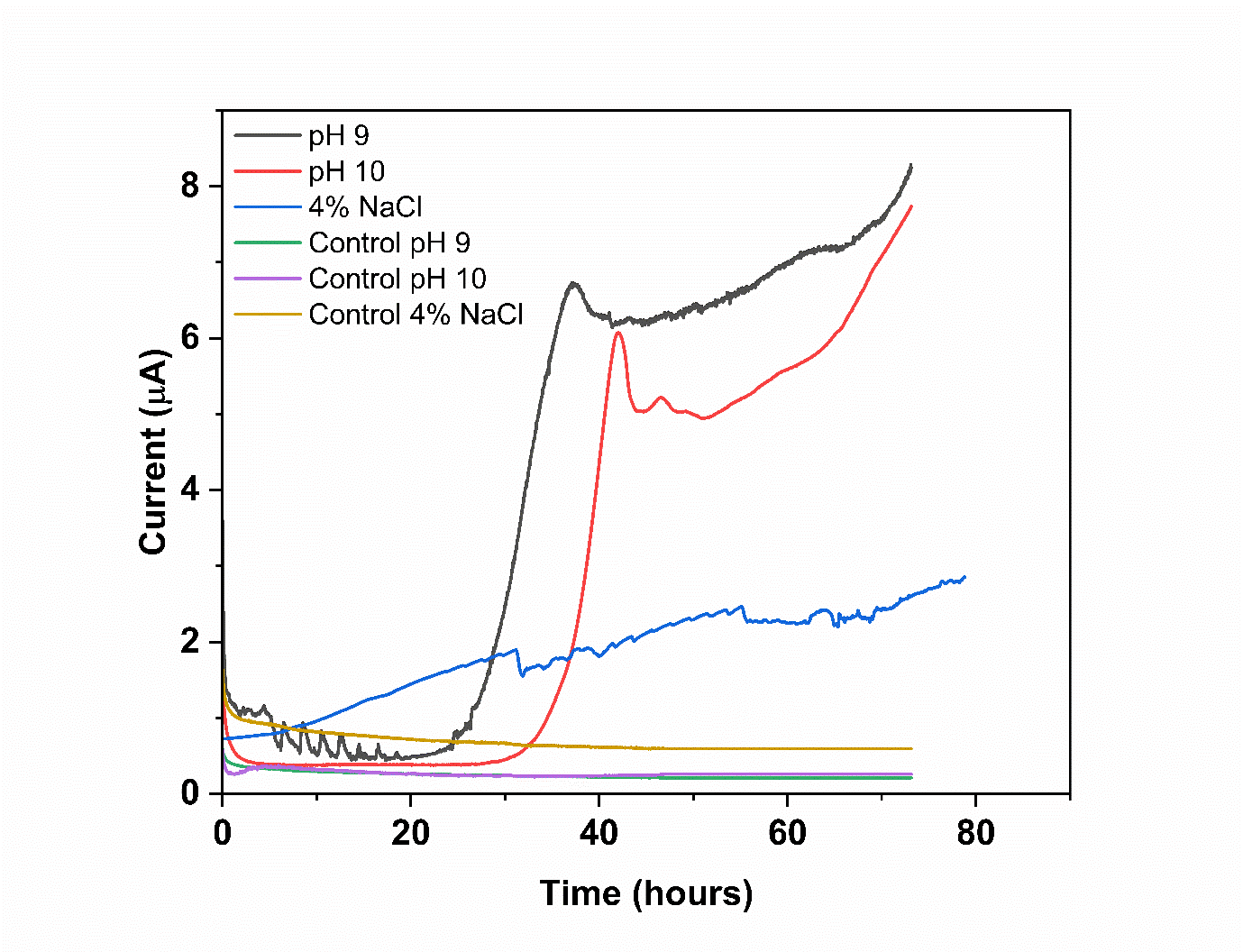

Figures S5: EET by *M. deferre* A1-JK in pH 9 and 10, and under halophilic conditions (4% salt, pH 7).

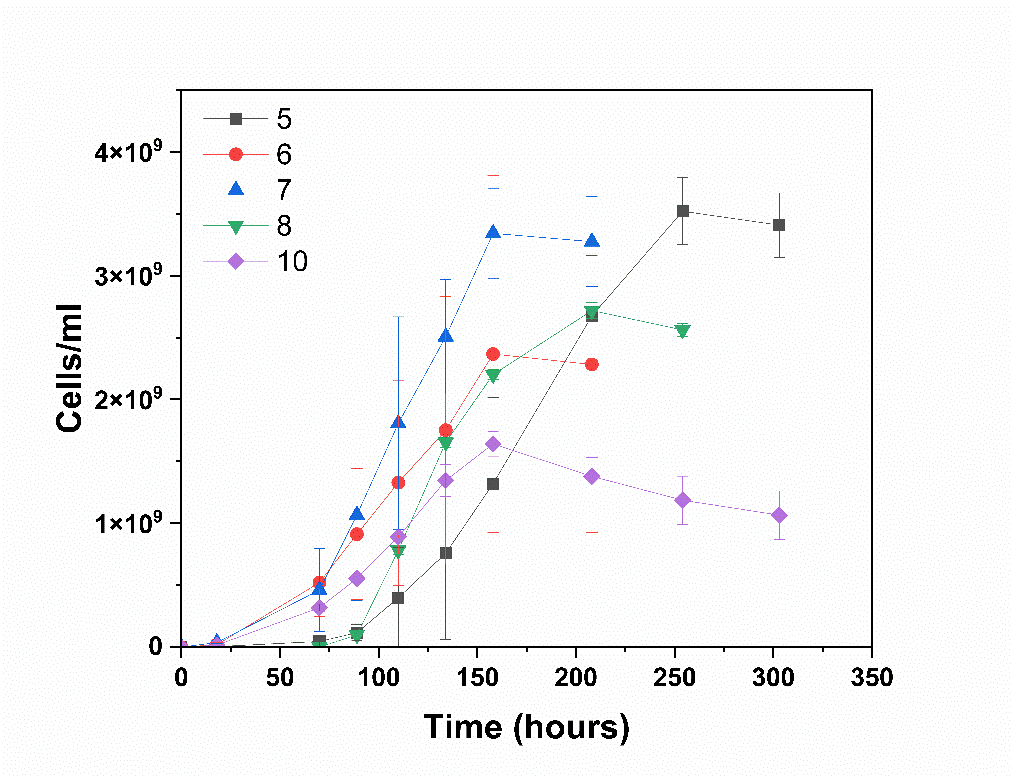

Figure S6: Growth curves of *M. deferre* A1-JK with a pH range from 5-10.

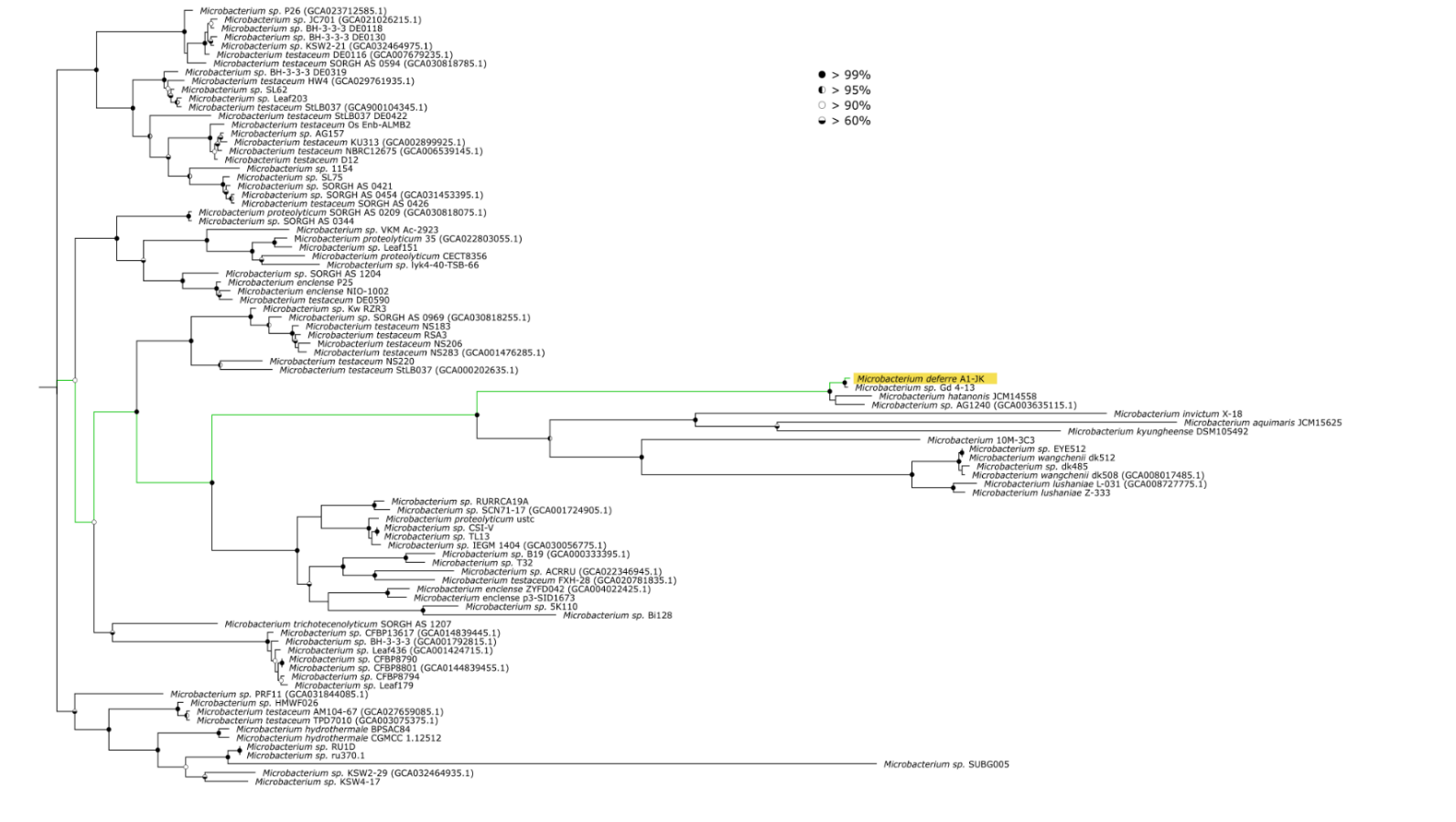

Figure S7: Phylogenetic tree of a selection of *Microbacterium* genomes available at NCBI. Yellow bar: *M.deferre* A1-JK, green lines: originating lineage of A1-JK

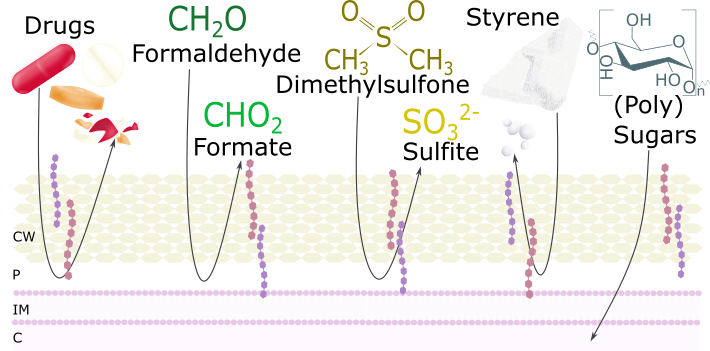

Figure S8: Selected gene-based metabolic capacities of *M. deferre* A1-JK. CW, cell wall; P, periplasm; IM, inner membrane; C, cytoplasm. Genomic analyses revealed the ability to degrade drugs like polystyrene, dimethylsulfone and styrene. (Poly)sugars can be broken down and assimilated for metabolic growth

Table S1: description of genomic and physiological characteristics of *M. deferre* A1-JK.
LB: lysogeny broth; NB: nutrient broth; R2A: Reasoner’s 2A broth; SBM: *Shewanella* basal medium. +: growth; ±: weak growth; -±: very weak growth; -: no growth, blank: untested.

| **Characteristics of A1-JK** |  | | |  | |  | |
| --- | --- | --- | --- | --- | --- | --- | --- |
| genome size | 3.46 Mb | | |  | |  | |
| G+C content | 69.32% | | |  | |  | |
| *growth at* |  | | |  | |  | |
| pH | **5** | **6** | **7** | | **8** | **9** | **10** |
|  | + | + | + | | + | + | + |
| Temperature (°C) | **4** | **15** | **18** | | **22** | **25** | **37,5** |
|  | ± | + | + | | + | + | -± |
| Salinity (% NaCl) | **1** | **2** | **3** | | **4** | **10** |  |
|  | + | + | + | | + | -± |  |
| Media | **LB** | **NB** | **R2A** | | **SBM** |  |  |
|  | + | + | + | | + |  |  |
| *growth on* | **24h** | | | **48h** | | **72h** | |
| 2-nitrophenyl-βD-galactopyranoside | - | | | - | | ± | |
| 4-nitrophenyl-βD-galactopyranoside | - | | | -± | | ± | |
| esculin | - | | | -± | | ± | |
| gelatine | - | | | - | | - | |
| L-arginine | - | | | - | | - | |
| L-lysine | - | | | - | | - | |
| L-ornithine | - | | | - | | - | |
| L-tryptophan |  | | | ± | |  | |
| trisodium citrate | - | | | + | |  | |
| urea | - | | | - | | - | |
| *production of* |  | | |  | |  | |
| acetoin |  | | | + | |  | |
| H_2_S | - | | | - | | - | |
| indole | - | | | - | |  | |
| NO_2_^-^ | - | | | - | |  | |
| N_2_ | ± | | | ± | |  | |
| *assimilation (A)/fermentation (F) of* | **A F** | | | **A F** | | **A F** | |
| amygdalin |  | ± | |  | + |  | + |
| adipic acid | - |  | | - |  | - |  |
| capric acid | - |  | | - |  | - |  |
| D-glucose | - | + | | + | + | + | + |
| malic acid | - |  | | - |  | - |  |
| D-mannitol | - | ± | | + | + | + | + |
| D-mannose | - |  | | ± |  | + |  |
| D-maltose | - |  | | ± |  | + |  |
| D-melibiose |  | -± | |  | -± |  | -± |
| D-sorbitol |  | - | |  | -± |  | -± |
| D-sucrose |  | ± | |  | + |  | + |
| gluconate | - |  | | ± |  | + |  |
| Inositol |  | - | |  | - |  | - |
| L-arabinose | - | + | | + | + | + | + |
| L-rhamnose |  | ± | |  | ± |  | ± |
| N-acetyl-glucosamine | - |  | | - |  | - |  |
| Phenylacetic acid | *-* |  | | *-* |  | *-* |  |

Table S2: Locus tags of the proteins, translated from the genome of *Microbacterium deferre* A1-JK, with e-values and bit scores of blastp, ran against a database of known EET protein sequences. Additionally, localization predictions and scores from psortb of *M. deferre* proteins, and the bacterial origin of the matched protein is shown.

| **Locustag** | **Protein** | **E-value** | **Bitscore** | **Cellular location** | **Score** | **Origin of database protein** |
| --- | --- | --- | --- | --- | --- | --- |
| A1_00037 | PhzE | 7,90E-06 | 462 | Cytoplasmic | 7,50 | *Pseudomonas aeruginosa* PAO1 |
| A1_00096 | ExtH | 1,06E-08 | 543 | Cytoplasmic | 9,97 | *Geobacter sulfurreducens* PCA |
| A1_00129 | LpdG | 1,55E-76 | 245 | Cytoplasmic | 9,97 | *Pseudomonas aeruginosa* |
| A1_00214 | PhzE | 4,51E-15 | 755 | Cytoplasmic | 9,97 | *Pseudomonas aeruginosa* PAO1 |
| A1_00269 | FccA | 9,01E-05 | 435 | Cytoplasmic | 7,50 | *Shewanella oneidensis* |
| A1_00292 | MenE | 4,73E-43 | 159 | Cytoplasmic | 9,97 | *Enterococcus faecalis* OG1RF |
| A1_00504 | DmkB | 1,30E-12 | 662 | Cytoplasmic | 9,97 | *Enterococcus faecalis* OG1RF |
| A1_00560 | FccA | 1,98E-22 | 994 | Cytoplasmic | 9,97 | *Shewanella oneidensis* |
| A1_00628 | Ndh3 | 1,75E-30 | 123 | Cytoplasmic membrane | 9,51 | *Enterococcus faecalis* OG1RF |
| A1_00730 | MenE | 6,52E-12 | 670 | Unknown | x | *Enterococcus faecalis* OG1RF |
| A1_00781 | LpdG | 2,34E-47 | 169 | Cytoplasmic | 9,97 | *Pseudomonas aeruginosa* |
| A1_00804 | FccA | 1,12E-32 | 131 | Cytoplasmic membrane | 8,78 | *Shewanella oneidensis* |
| A1_00812 | RibD | 2,36E-70 | 222 | Cytoplasmic | 7,50 | *Escherichia coli* K-12 |
| A1_00813 | RibE1 | 2,47E-27 | 102 | Cytoplasmic | 9,97 | *Aeromonas hydrophila* 4AK4 |
| A1_00814 | RibBA | 6,97E-68 | 218 | Cytoplasmic | 7,50 | *Shewanella oneidensis* MR-1 |
| A1_00815 | RibE2 | 1,84E-27 | 100 | Unknown | x | *Aeromonas hydrophila* |
| A1_00836 | LpdG | 1,88E-36 | 138 | Cytoplasmic | 9,67 | *Pseudomonas aeruginosa* |
| A1_01299 | PhzD | 1,76E-08 | 512 | Cytoplasmic | 7,50 | *Pseudomonas aeruginosa* PAO1 |
| A1_01320 | MenE | 4,34E-23 | 100 | Cytoplasmic | 9,97 | *Enterococcus faecalis* OG1RF |
| A1_01353 | LpdG | 1,43E-09 | 589 | Cytoplasmic | 9,97 | *Pseudomonas aeruginosa* |
| A1_01395 | MenE | 2,03E-13 | 709 | Cytoplasmic | 9,97 | *Enterococcus faecalis* OG1RF |
| A1_01404 | MenE | 4,58E-37 | 141 | Cytoplasmic | 9,97 | *Enterococcus faecalis* OG1RF |
| A1_01430 | MenB | 1,51E-27 | 105 | Cytoplasmic | 7,50 | *Enterococcus faecalis* OG1RF |
| A1_01457 | MenB | 1,79E-16 | 755 | Unknown | x | *Enterococcus faecalis* OG1RF |
| A1_01465 | FmnA | 2,24E-09 | 551 | Cytoplasmic membrane | 10,00 | *Listeria monocytogenes* |
| A1_01472 | MenB | 3,26E-11 | 620 | Cytoplasmic | 7,50 | *Enterococcus faecalis* OG1RF |
| A1_01539 | MenE | 1,04E-73 | 239 | Cytoplasmic | 9,97 | *Enterococcus faecalis* OG1RF |
| A1_01580 | YeeO | 5,98E-14 | 720 | Cytoplasmic membrane | 10,00 | *Escherichia coli* K-12 |
| A1_01655 | ExtH | 7,56E-10 | 574 | Cytoplasmic | 9,97 | *Geobacter sulfurreducens* PCA |
| A1_01752 | PhzE | 6,28E-16 | 743 | Cytoplasmic | 9,64 | *Pseudomonas aeruginosa* PAO1 |
| A1_01950 | FmnA | 7,07E-08 | 508 | Cytoplasmic membrane | 10,00 | *Listeria monocytogenes* |
| A1_02039 | LpdG | 4,72E-11 | 628 | Cytoplasmic | 9,97 | *Pseudomonas aeruginosa* |
| A1_02057 | MenE | 5,30E-38 | 144 | Cytoplasmic | 9,97 | *Enterococcus faecalis* OG1RF |
| A1_02167 | MenC | 3,23E-07 | 497 | Cytoplasmic | 7,50 | *Escherichia coli* K-12 |
| A1_02170 | MenB | 4,66E-84 | 253 | Cytoplasmic | 7,50 | *Enterococcus faecalis* OG1RF |
| A1_02171 | MenE | 1,48E-34 | 131 | Unknown (extracellular or cellwall) | 5,15; 4,55 | *Enterococcus faecalis* OG1RF |
| A1_02172 | DmkA | 1,76E-11 | 624 | Cytoplasmic membrane | 10,00 | *Enterococcus faecalis* OG1RF |
| A1_02180 | PhzE | 1,24E-11 | 647 | Cytoplasmic | 9,97 | *Pseudomonas aeruginosa* PAO1 |
| A1_02183 | DmkB | 3,63E-25 | 102 | Cytoplasmic | 9,97 | *Enterococcus faecalis* OG1RF |
| A1_02340 | MenE | 6,58E-21 | 947 | Cytoplasmic | 9,97 | *Enterococcus faecalis* OG1RF |
| A1_02377 | MenE | 1,05E-11 | 674 | Cytoplasmic membrane | 9,60 | *Enterococcus faecalis* OG1RF |
| A1_02719 | DmkB | 2,86E-18 | 832 | Cytoplasmic | 9,97 | *Enterococcus faecalis* OG1RF |
| A1_02726 | MenE | 6,11E-18 | 855 | Cytoplasmic | 9,97 | *Enterococcus faecalis* OG1RF |
| A1_02735 | LpdG | 1,34E-97 | 300 | Cytoplasmic | 9,97 | *Pseudomonas aeruginosa* |
| A1_02870 | PhzE | 0,00E+00 | 525 | Cytoplasmic | 9,97 | *Pseudomonas aeruginosa* PAO1 |
| A1_02934 | PhzG | 1,28E-34 | 121 | Cytoplasmic membrane | 8,16 | *Pseudomonas fluorescens* |
| A1_03020 | LpdG | 1,70E-10 | 612 | Cytoplasmic | 9,97 | *Pseudomonas aeruginosa* |
| FLEET* | EetA | Missing | nd | nd | nd | *L. monocytogenes/E. faecalis* |
| FLEET* | EetB | Missing | nd | nd | nd | *L. monocytogenes/E. faecalis* |
| FLEET* | PplA | Missing | nd | nd | nd | *Listeria monocytogenes* |

*Amino acid sequences part of the flavin-EET (FLEET) pathway, common for Gram-positive microbes, these were all missing in the A1-JK translated genome.
